# Research grade marijuana supplied by the National Institute on Drug Abuse is genetically divergent from commercially available *Cannabis*

**DOI:** 10.1101/592725

**Authors:** Anna L. Schwabe, Connor J. Hansen, Richard M. Hyslop, Mitchell E. McGlaughlin

## Abstract

Public comfort with *Cannabis* (marijuana and hemp) has recently increased, resulting in previously strict *Cannabis* regulations now allowing hemp cultivation, medical use, and in some states, recreational consumption. There is a growing interest in the potential medical benefits of the various chemical constituents produced by the *Cannabis* plant. Currently, the University of Mississippi, funded through the National Institutes of Health/National Institute on Drug Abuse (NIH/NIDA), is the sole Drug Enforcement Agency (DEA) licensed facility to cultivate *Cannabis* for research purposes. Hence, most federally funded research where participants consume *Cannabis* for medicinal purposes relies on NIDA-supplied product. Previous research found that cannabinoid levels in research grade marijuana supplied by NIDA did not align with commercially available *Cannabis* from Colorado, Washington and California. Given NIDA chemotypes were misaligned with commercial *Cannabis*, we sought to investigate where NIDA’s research grade marijuana falls on the genetic spectrum of *Cannabis* groups. NIDA research grade marijuana was found to genetically group with Hemp samples along with a small subset of commercial drug-type *Cannabis*. A majority of commercially available drug-type *Cannabis* was genetically very distinct from NIDA samples. These results suggest that subjects consuming NIDA research grade marijuana may experience different effects than average consumers.

## Introduction

Humans have a long history with *Cannabis sativa* (marijuana and hemp), with evidence of cultivation dating back as far as 10,000 years ago^1^. The World Health Organization proclaims *Cannabis* as the most widely cultivated, trafficked and abused illicit drug, and reports over half of worldwide drug seizures are of *Cannabis*^2^. Phytochemicals of interest in *Cannabis* are primarily Δ^9^-tetrahydrocannabinolic acid (THCA) and cannabidiolic acid (CBDA), both of which require a decarboxylation conversion to the biologically active forms, THC and CBD, respectively. The United States is currently experiencing drastic changes in patterns of *Cannabis* use associated with widespread relaxation of laws that previously limited both medical and recreational marijuana consumption^3^ and hemp cultivation. This has led to a need for extensive research into the basic biology and taxonomy of *Cannabis sativa*^4–8^, and the possible benefits and threats from *Cannabis* consumption^3,9^.

Although *Cannabis sativa* is the only described species in the genus *Cannabis* (Cannabaceae), there are several commonly described subcategories of *Cannabis* that are widely recognized. There are two primary *Cannabis* usage groups, which are well supported by genetic analyses^7,10–12^: ***Hemp*** is defined by a lack of THC (< 0.3% THC in the U.S.), and ***marijuana*** or ***drug-types*** have moderate to high THC concentrations (> 0.3% THC in the U.S.). Hemp-type *Cannabis* tends to have higher concentrations of CBD than drug-types^13^. Drug-type *Cannabis* usually contains > 12% THC and averages ~ 10-23% THC in commercially available dispensaries^14–16^. Within the two major usage groups, *Cannabis* can be further divided into varietals, which are referred to as strains. The drug-type strains are commonly categorized further: ***Sativa*** strains reportedly have uplifting and more psychedelic effects, ***Indica*** strains reportedly have more relaxing and sedative effects, and ***Hybrid*** strains, which result from breeding Sativa and Indica strains, have a spectrum of intermediate effects. There is extensive debate among experts surrounding the appropriate taxonomic treatment of *Cannabis* groups, which is confounded by colloquial usage of these terms versus what researchers suggest is more appropriate nomenclature^5,17–24^ Commercially available drug-type strains for medical or recreational consumption are labeled with a strain name, as well as the levels of THC and often CBD as a percent of the dry weight. Genetic analyses have not shown clear and consistent differentiation among the three commonly described drug-type strains^7,10^, but both the recreational and medical *Cannabis* communities maintain there are distinct differences in effects between Sativa and Indica strains^25–27^.

*Cannabis* has been federally controlled since 1937, many states now allow regulated medical (33 states and the District of Columbia) and recreational use (10 states and the District of Columbia)^28^. There were > 3.5 million registered medical marijuana patients reported as of May 2018^29^. However, the United States Drug Enforcement Agency (DEA) lists *Cannabis sativa* as a Schedule 1 substance^30^, and as such, research on all aspects of this plant has been limited. U.S. Surgeon General Jerome Adams recently expressed concern that the current scheduling in the most restrictive category is inhibiting research on *Cannabis* as a potentially therapeutic plant^31^. A Schedule 1 substance is described as a drug with no accepted medical use and a high potential for abuse^30^. The University of Mississippi, funded through the National Institutes of Health/National Institute on Drug Abuse (NIH/NIDA), currently holds the single license issued by the DEA for the cultivation of *Cannabis* for research purposes^32^. As such, NIDA serves as the sole legal provider of *Cannabis* for federally funded medical research in the United States. Bulk research grade marijuana supplied by NIDA is characterized by the level of THC and CBD. They offer *Cannabis* for research with four levels of THC: ***low***(< 1%), ***medium***(1-5 %), ***high***(5-10 %) and ***very high***(>10%), with the additional option of four levels of CBD: ***low***(< 1%), ***medium***(1-5%), ***high***(5-10%) and ***very high***(> 10%).

The National Institute on Drug Abuse funds a wide range of research on drug-type *Cannabis*, including long and short-term effects on behavior, pain, mental illness, brain development, use and abuse, and impacts of policy changes related to marijuana^33,34^ Additionally, the NIH provides support for researching cannabinoids as separate constituents. Funding for CBD related research is reported as $36M (2015 - 2017) and projected to be $36M for 2018 - 2019^35^, while cannabinoid related research is reported as $366M from 2015 - 2017 and projected to be $292M for 2018 - 2019^36^.

Recent research has documented that NIDA-provided *Cannabis* has distinctly different cannabinoid profiles than commercially available *Cannabis*^14^ Specifically, Vergara et al. (2017) found that NIDA samples contained only 27% of the amount of THC and 48% of CBD levels of commercially available *Cannabis*. The substantial chemical differences between NIDA and commercially available *Cannabis* raises significant questions about whether research conducted with federal *Cannabis* is indicative of the experience consumers are having.

Medical research on *Cannabis* primarily focuses on THC and CBD^3,9,35–40^, but there are hundreds of other chemical constituents in *Cannabis*^41^, including cannabinoids and terpenes, which have largely been ignored^9^. There is evidence to suggest that chemical constituents in various combinations and abundances work in concert to create the suite of physiological effects reported^9^. The chemical makeup of each variant of *Cannabis* is influenced by the genetic makeup as well as environmental conditions. Given that previous research has determined the cannabinoid levels of research grade marijuana from NIDA is significantly different from commercially available *Cannabis*^14^, genetic investigations are warranted to determine if NIDA *Cannabis* is genetical distinct from other sources. In the current study we investigated the genetic relationship of NIDA provided *Cannabis* to commercially available drug-type strains, as well as feral and cultivated hemp. Ten variable nuclear microsatellite regions were used to examine genetic differentiation among our samples. Sampling included NIDA (High THC and High THC/CBD), high THC drug-type, low THC/high CBD drug-type, wild growing hemp (presumed escapees from cultivation), and commercial hemp. This study aimed to investigate where research grade marijuana supplied by NIDA falls on the genetic spectrum of *Cannabis* groups.

## Results

Our analyses examined the genetic differentiation and structure of samples from six groups (Supplemental Table 1). 1) **NIDA** – research grade marijuana samples obtained from NIDA classified as High THC or High THC/CBD; 2) **Hemp** – *Cannabis* obtained from hemp cultivators and feral collected hemp; 3) **High CBD** – drug-type *Cannabis* with relatively high levels of CBD and low levels of THC; and commercially available drug-type *Cannabis* described as 4) **Sativa**, 5) **Hybrid**, or 6) **Indica** strains. Analyses were also performed on samples at the individual level to control for biases that might arise due to the potential artificial nature of named groups and varying group sample sizes.

### Genetic Differentiation

Pairwise genetic differentiation (Fst and Nei’s D) calculated in GENALEX ver. 6.4.1 (Peakall & Smouse 2006, Peakall & Smouse 2012) found the highest level of divergence between hemp and high CBD drug-type strains (Fst = 0.215) and between hemp and Sativa drug-type strains (Nei’s D = 0.614) (Table 1). The least divergence was observed among the drug-type strains (Fst = 0.023-0.04; Nei’s D = 0.066-0.109).

**Table 1.** Pairwise Fst values (below the diagonal) and Nei’s D (above the diagonal) for major *Cannabis* groups.

|  | NIDA | Hemp | High CBD | Sativa | Hybrid | Indica |
| --- | --- | --- | --- | --- | --- | --- |
| NIDA |  | 0.519 | 0.527 | 0.553 | 0.480 | 0.441 |
| Hemp | 0.120 |  | 0.489 | 0.614 | 0.585 | 0.459 |
| High CBD | 0.166 | 0.215 |  | 0.329 | 0.310 | 0.281 |
| Sativa | 0.114 | 0.160 | 0.137 |  | 0.098 | 0.109 |
| Hybrid | 0.117 | 0.149 | 0.135 | 0.040 |  | 0.066 |
| Indica | 0.078 | 0.124 | 0.121 | 0.035 | 0.023 |  |

### Clustering Analysis

Principal Coordinate Analysis (PCoA) was conducted in GENALEX and plotted in R Studio with the ggplot package^42^ with 95% confidence interval ellipses around the major groups (Figure 1). No confidence intervals were drawn for NIDA (n = 2) or High CBD (n = 3) due to small sample size. Coordinate 1 explains 13.26% of the genetic variation and an additional 11.39% of the genetic variation is explained by coordinate 2. The drug-type strains (Indica, Sativa, Hybrid, and High CBD) all occupy the same character space. There is clear separation of hemp samples from the drug-types, with NIDA samples clustering within the hemp confidence interval.

**Figure 1:**
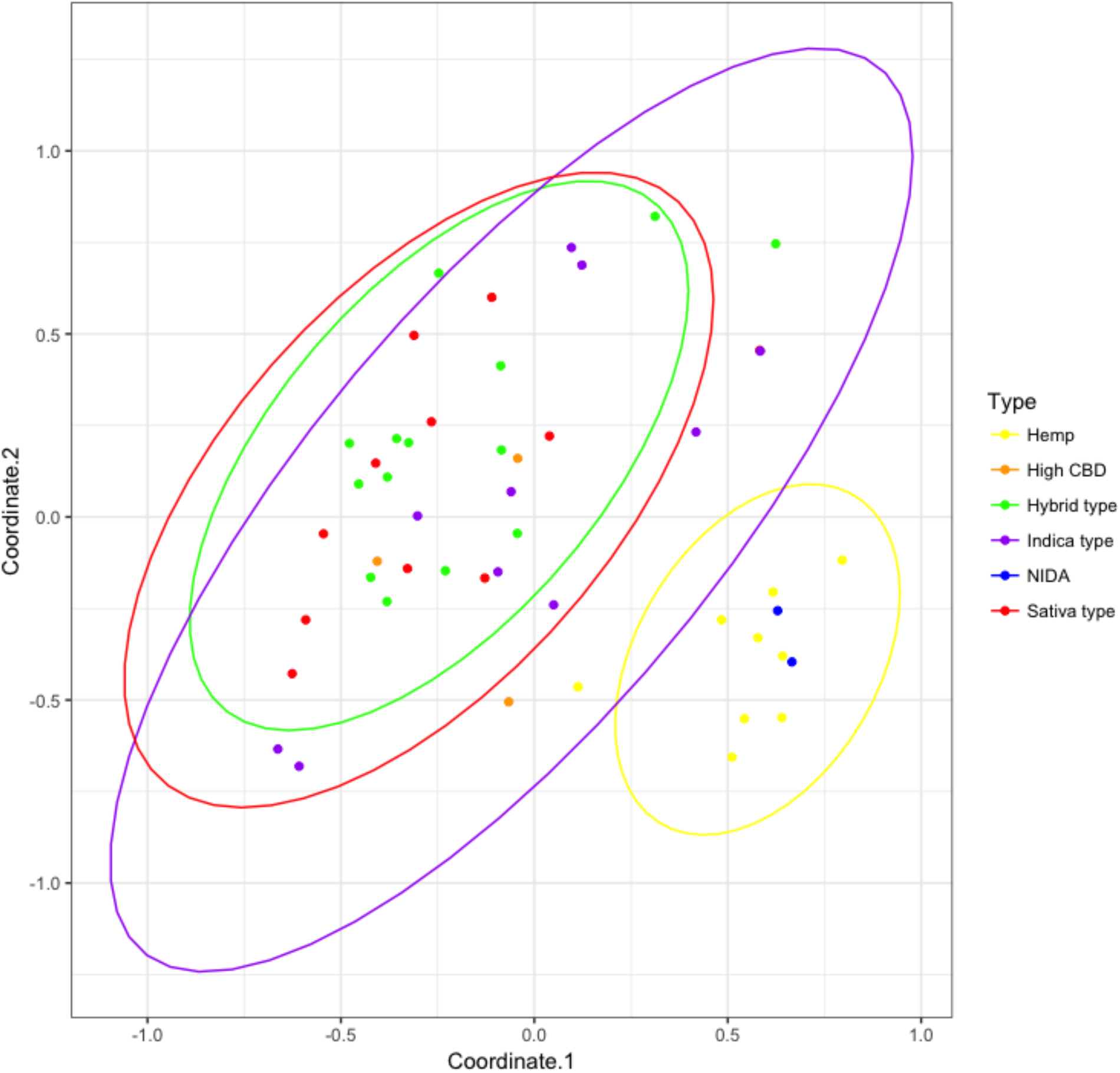
Principal Coordinates Analysis with 95% confidence intervals around the major groups (hemp = yellow, NIDA = blue, High CBD = orange, Sativa = red, Hybrid = green, Indica = purple). Approximately 25% of the genetic variation in these groups is shown (coordinate 1= 13.26% and coordinate 2 = 11.39%). No confidence intervals were drawn for NIDA or High CBD samples due to the small sample size (n = 2 and n = 3, respectively).

PC-Ord version 6^43^ was used to generate a dendrogram with Ward’s method and Euclidean Genetic distance parameters based on pairwise genetic distance values generated in GENALEX (Figure 2). The initial branching split the samples into two clusters, A and B. Cluster A contains all but one hemp sample (88%), as well as the NIDA samples (100%) and two drug-type samples (5%). Cluster B contains the remaining drug-type samples (95%) and one hemp sample (12%).

**Figure 2:**
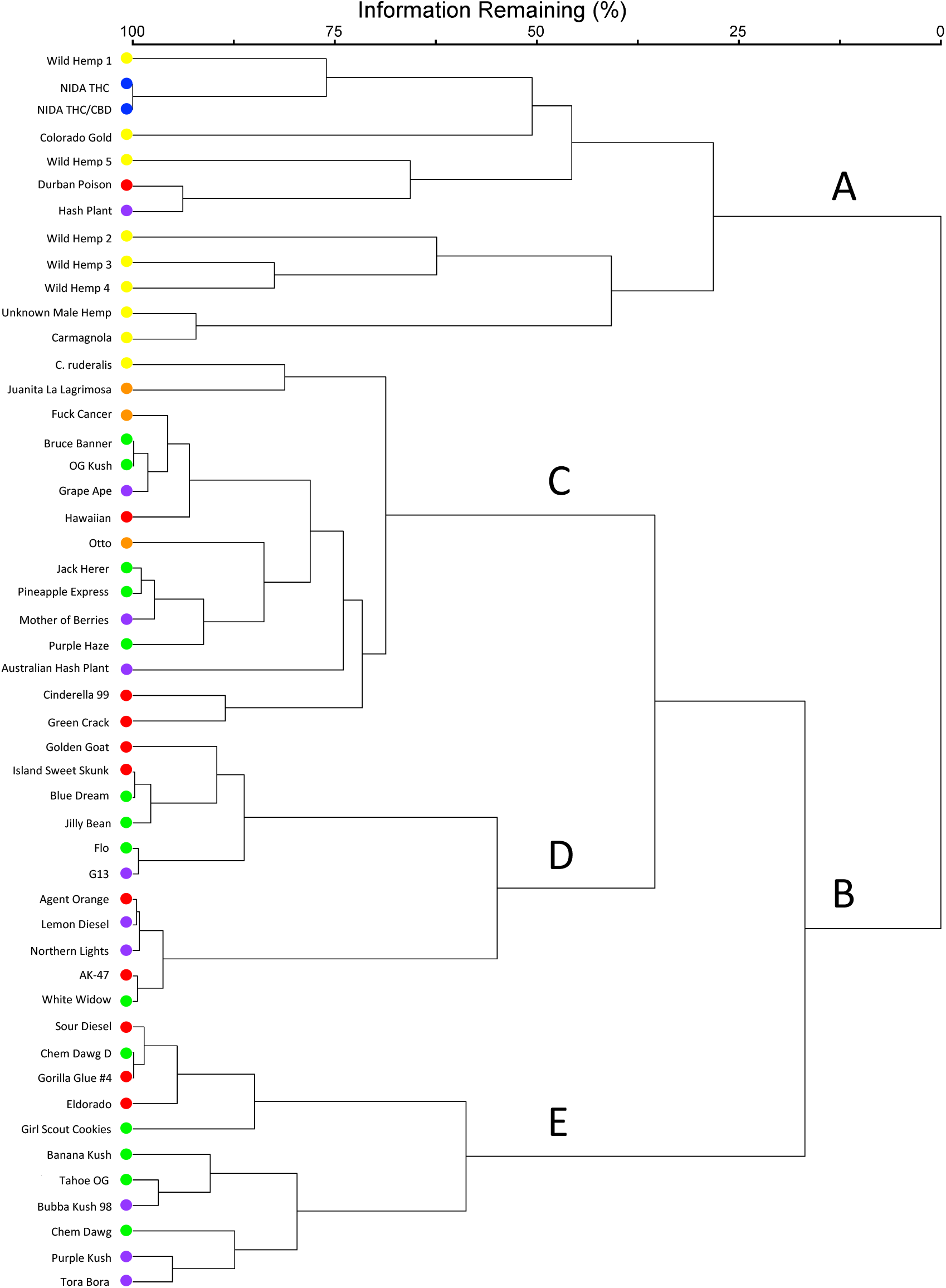
PC-Ord group linkage dendrogram. Samples are color-coded (Hemp = yellow, NIDA = blue, High CBD = orange, Sativa = red, Hybrid = green, Indica = purple). Cluster B further branches into three clusters (C, D, and E), where Sativa, Hybrid and Indica drug type strains are dispersed throughout.

STRUCTURE ver. 2.4.2^44^ was used to examine sample assignment to genetic groups while allowing admixture. The appropriate number of STRUCTURE groups was validated using STRUCTURE HARVESTER^45^, which had high support for two genetic groups (K = 2, ΔK = 67.68) and weak support for three genetic groups (K = 2, ΔK = 4.48) (Supplemental Figure 1). Additionally, MavericK 1.0.5^46^ was used to independently test group assignments, which also had strong support for two genetic groups (K = 2, probability 0.901) and weaker support for three genetic groups (K = 3, probability 0.097) (Supplemental Figure 2), with the sample assignments matching STRUCTURE (Supplemental Figure 3). The two genetic group STRUCTURE analyses (Figure 3) show consistent differentiation between hemp and drug-type strains. All hemp samples were assigned to genetic group 1 (yellow) with a proportion of inferred ancestry (Q) greater than 0.82 (hemp mean group 1, Q = 0.94). Drug-type samples showed some admixture with the majority of the genetic signal of 31 samples (82%) being assigned to genetic group 2 (green; drug-type mean group 2, Q = 0.72). NIDA samples were assigned to genetic group 1 (NIDA mean group 1, Q = 0.97), demonstrating a strong association with hemp. Although not strongly supported, the three genetic group analysis shows some additional genetic structure among drug-type strains.

**Figure 3:**
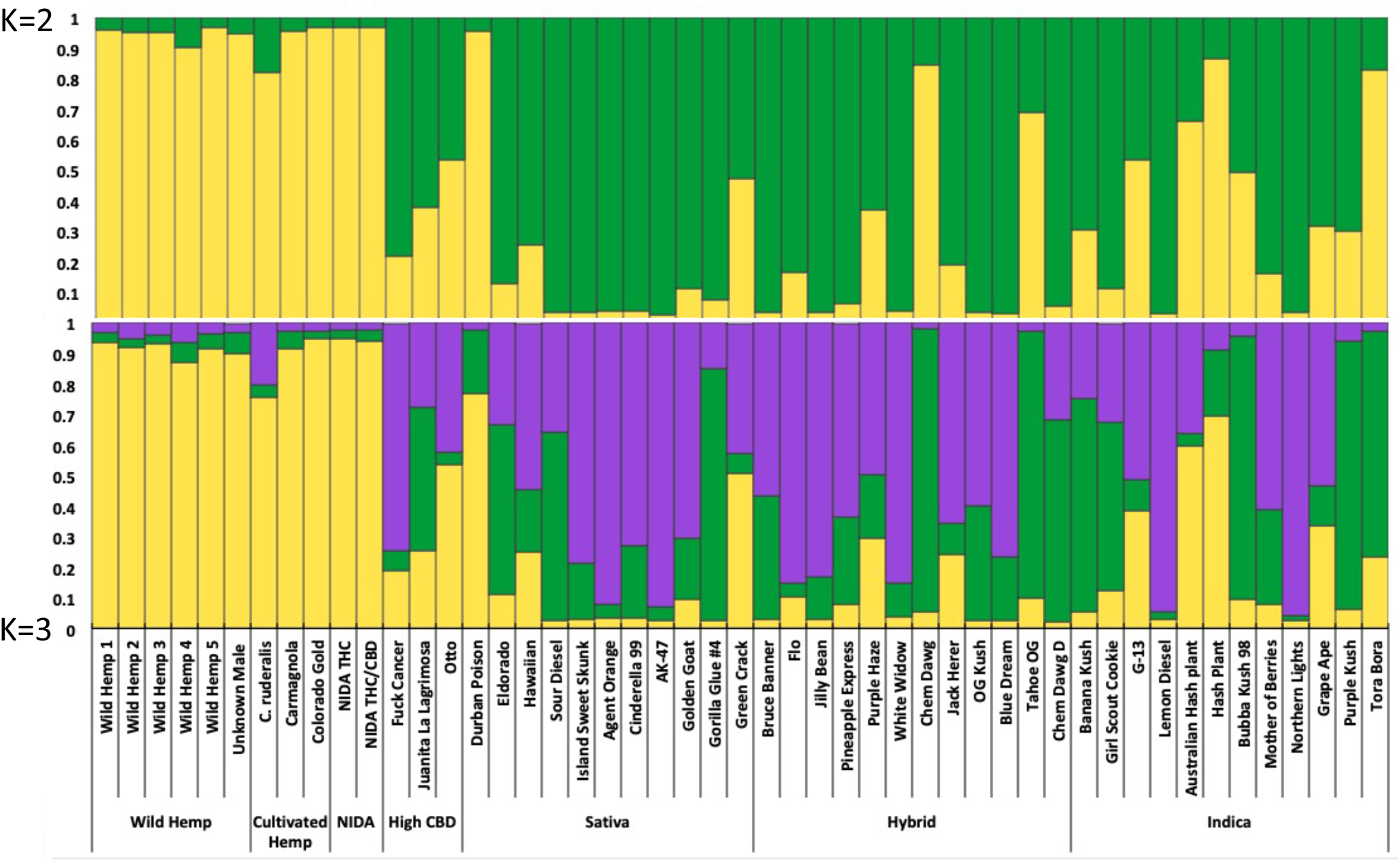
Bayesian clustering analysis from STRUCTURE with the proportion of inferred ancestry for two genetic groups (K = 2, top), and for three genetic groups (K = 3, bottom). Each individual is represented as a single bar in the graph.

EDENetwork ver. 2.18^47^ was used to generate a web of genetic relationship based on pairwise linkages (Figure 4). The automatically selected percolation threshold was 8.1 (Figure 4A), although not all individuals were connected at this level. The threshold was raised iteratively to connect more divergent samples and explore larger patterns of genetic relationships. The two NIDA samples were united at a threshold of 8.5 (Figure 4B). When the threshold was raised to 13.7 (Figure 4C) the NIDA samples became connected to the network via the drug-type sample Eldorado. At a threshold level of 16.9 (Figure 4D) all samples in the dataset are included in the relationship network.

**Figure 4:**
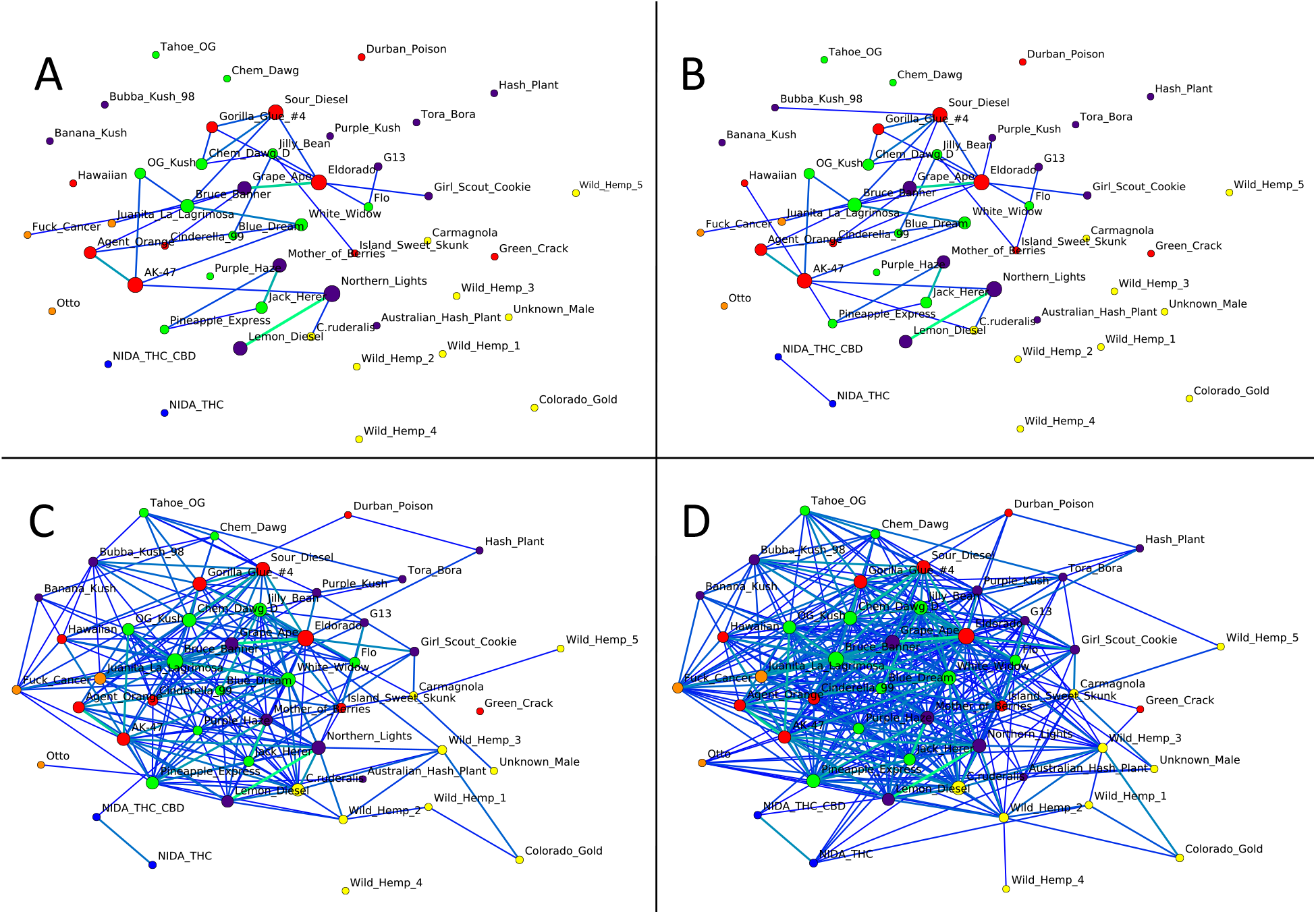
EDENetworks genetic relationship network with incrementally decreasing stringency of required genetic relatedness among samples in the data set. (A) Threshold 8.1: the percolation threshold determined by the analysis. (B) Threshold 8.5: the threshold required to connect NIDA samples to each other, but not to any other samples in the dataset. (C) Threshold 13.7: the threshold necessary to connect the NIDA sample to the larger network with the connection via the drug-type strain Eldorado. (D) Threshold 16.9: the required threshold to connect all samples in the network. Nodes are colored to indicate group designation (Hemp = yellow, NIDA = blue, High CBD = orange, Sativa = red, Hybrid = green, Indica = purple). Node size is proportionate to the number of connections to that individual within the network. Lines thinner and lighter in color indicate weak genetic relationships, while thicker darker lines indicate stronger relationships.

## Discussion

The purpose of this study was to examine the genetic relationship of *Cannabis* samples from the National Institute on Drug Abuse (NIDA) to hemp and drug-type samples. Our results clearly demonstrate that NIDA *Cannabis* samples are substantially different from most commercially available drug-type strains, sharing a genetic affinity with hemp samples in most analyses. Previous research has found that medical and recreational *Cannabis* from California, Colorado, and Washington differs significantly in cannabinoid levels from the research grade marijuana supplied by NIDA^14^ Our genetic investigation adds to this previous research, indicating that the genetic makeup of NIDA *Cannabis* is also distinctive from commercially available medical and recreational *Cannabis*.

The genetic data collected in this study indicate that two major genetic groups exist within *Cannabis sativa*. The first group contained a majority of hemp (88 - 100%, depending on analysis) and both NIDA samples (100%), while the second group contained a majority of drug-type samples (82 - 95%). These results contribute to the growing consensus that hemp and drug-type *Cannabis* can be consistently differentiated^7,10–12,48–51^. To our knowledge, this is the first genetic study to include research grade marijuana from NIDA, and its placement with hemp samples was unexpected. However, it is important to note that some drug-type samples (e.g. Durban Poison, Figure 2 & 3) are also placed in the hemp group. Although the sample size of NIDA samples could impact their placement in group-based analyses such as genetic distances (Table 1), all other analyses were carried out at an individual level (Figures 1 – 4) to avoid this issue.

According to the University of Mississippi National Center for Natural Products Research (NCNPR), which produces research grade marijuana for NIDA, the first experimental plots of *Cannabis* were planted in 1968 with seeds from “Mexico, Panama, Southeast Asia, Korea, India, Afghanistan, Iran, Pakistan, and Lebanon”^52,53^. Over the next decade, cultivation techniques were standardized, with over 100 varieties planted in 1976^52^. Between the late 1970’s and today, the University of Mississippi has continued to be the sole producer of research grade marijuana for NIDA, and it has refined cultivation techniques and extraction procedures, particularly for THC and CBD^54^. The program does not provide variety or strain information when filling *Cannabis* orders, so it is unclear what is currently grown by NCNPR for federally funded marijuana research. The NCNPR director recently stated that “The marijuana project currently stocks 27 plant varieties with different cannabinoid profiles, various CBG potencies, and a wide range of THC levels”^53^. However, the NCNPR website states that only three *Cannabis* varieties were grown in 2014^52^. Our data suggest that the NIDA *Cannabis* analyzed in this study was sourced from a single strain or two very closely related strains within the NCNPR stock. Without additional information about NCNPR *Cannabis* production, it is difficult to know how many strains are being used in research.

This study indicates the need for additional research and refinement of our understanding of *Cannabis* genetic structure and how those differences might impact *Cannabis* consumers. Although medicinal research on *Cannabis* has predominantly focused on THC and CBD^3,9,35–40^, it is becoming apparent that other chemical constituents in various combinations and abundances likely have important effects^9^. If researchers are solely interested in the effects of THC and CBD at known concentrations, then NIDA *Cannabis* could serve as a representative source, although in these cases, isolates of these molecules may be more appropriate. However, given the genetic distinction between NIDA and commercially available *Cannabis*, patients in federally funded *Cannabis* research are likely experiencing effects that are specific to the plant material provided by NIDA. As the interest for medical *Cannabis* increases, it is important that research examining the threats and benefits of *Cannabis* use accurately reflect the experiences of the general public.

Given the rapidly changing landscape of *Cannabis* regulations and consumption^28^, it is not surprising that commercially available *Cannabis* contains a diversity of genetic types.

Commercially available *Cannabis* has come to market through non-traditional means leading to many inconsistencies. We have previously documented^55^ that there is substantial genetic divergence among samples within named strains, which only exacerbates questions about the impacts of *Cannabis* consumption. These results emphasize the need to increase consistency within the *Cannabis* marketplace, and the need for research grade *Cannabis* to accurately represent what is accessible to consumers.

In conclusion, this study highlights the genetic difference between research grade marijuana provided by NIDA and commercial *Cannabis* available to medical and recreational users. This finding reveals that research conducted with NIDA *Cannabis* may not be indicative of the effects that consumers are experiencing. Additionally, research has demonstrated that *Cannabis* distributed by NIDA has lower levels of the principal medicinal cannabinoids (THC and CBD) and higher levels of degradation byproducts of cannabinoids (cannabinol, CBN)^14^ Taken together, these results demonstrate the need for there to be greater diversity of *Cannabis* available for medical research and that the genetic provenance of those samples to be established to fully understand the implications of results.

## Methods

A total of 49 *Cannabis* samples were used in this research (Supplemental Table 1), including: wild hemp (5), cultivated hemp (4), NIDA strains (2), high CBD drug-type strains (3), and drug-types strains (35). Drug-type strains were further subdivided into three commonly used categories: Sativa (11), Hybrid (14), and Indica (10) based on information available online^27,56^. The drug-type strains were randomly chosen from a much larger pool of samples. Duplicate accessions within strains were not included.

DNA was extracted using a modified CTAB extraction protocol^57^ with 0.035-0.100 grams of dried flower tissue per extraction. Ten variable microsatellite loci developed by Schwabe and McGlaughlin^55^ were used in this study following their previously described procedures.

GENALEX ver. 6.4.1^59,60^ was used to calculate pairwise genetic differentiation (F_ST_) and Nei’s genetic distance (D) between each of the six groups. PCoA eigenvalues calculated in GENALEX were used to plot the PCoA in RStudio with the ggplot package^42,61^ with 95% confidence interval ellipses. GENALEX was also used to generate a pairwise genetic distance square matrix which was then used to generate a hierarchical cluster analysis dendrogram with Ward’s method and Euclidean Genetic distance parameters in PC-ORD^43^.

Genotypes were analyzed using the Bayesian cluster analysis program STRUCTURE ver. 2.4.2 44 Burn-in and run-lengths of 50,000 generations were used with ten independent replicates for each STRUCTURE analysis. The number of genetic groups for the data set was determined by STRUCTURE HARVESTER^45^, which implements the Evanno et al. method^62^.

Maverick v1.0.5^46^ was used as an additional verification of Bayesian clustering analysis using thermodynamic integration to determine the appropriate number of genetic groups. The following parameters were used: admixture parameter (alpha) of 0.03 with a standard deviation (alphaPropSD) of 0.008, 10 replicates (mainRepeats), 1,000 Burn-in iterations (mainBurnin), 5,000 sample iterations (mainRepeats), 100 TI rungs (thermodynamicRungs), 500 TI Burn-in iterations (thermodynamicBurnin), and 1,000 TI iterations (thermodynamicSamples).

EDENetworks ver. 2.18^47^ was used to construct a web of genetic relationships using the Linear Manhattan distance measure. An auxiliary data file was imported to maintain the spatial coordinates and to color individuals by group assignment. The automatic percolation threshold was first derived as 8.1. Networks were generated for subsequent iterative threshold intervals of 0.5. Increasing the threshold lowers the stringency for genetic relationships, and as the threshold increases, more relationships are formed in the network. EDENetworks diagrams were constructed for the percolation threshold of 8.1, 8.5, 13.7 and 16.9. These are the values that: connect NIDA samples to each other, but not to any other samples in the dataset (8.5), connect a single NIDA sample to the larger network (13.7), and finally connect all samples in the network (16.9). The size of each node is proportionate to the number of relationship connections to other members in the network. The line color and width indicated the strength of the relationship between two individuals-lighter thicker lines indicate stronger genetic relationships, while the darker thinner lines indicate weaker genetic relationships.

## Data Availability

The scored microsatellite data set analyzed in this study is provided as supplementary material (Supplemental Table 2).

## Acknowledgements

The National Institute on Drug Abuse provided the Research Grade *Cannabis* samples from which DNA used in this study was extracted. We thank Matt Kahl and Caren Kershner for providing hemp samples for this project, Melissa Islam, Associate Director of Biodiversity Research at the Denver Botanic Gardens for access to wild collected hemp herbarium specimens (Kathryn Kalmbach Herbarium), and the Cannabis Genome Research Initiative for the sample of *Cannabis ruderalis*. Funding for this project was provided through research grants awarded to A. Schwabe by the University of Northern Colorado Graduate Student Association and the University of Northern Colorado College of Natural and Heath Sciences, and the McGlaughlin Lab, School of Biological Sciences, University of Northern Colorado.

## Author Contributions

A.S conceived the project, collected samples, conducted DNA extractions, designed and optimized microsatellite primers, compiled and analyzed data, and drafted manuscript content; C.H conducted DNA extractions, compiled and analyzed data, and prepared the first draft of the manuscript; R.M.H provided DNA from NIDA samples; M.E.M directed the project, provided some funding, contributed statistical analysis and manuscript revisions; all authors contributed to manuscript preparation.

