## Supplemental Figures for "Research grade marijuana supplied by the National Institute on Drug Abuse is genetically divergent from commercially available *Cannabis*"

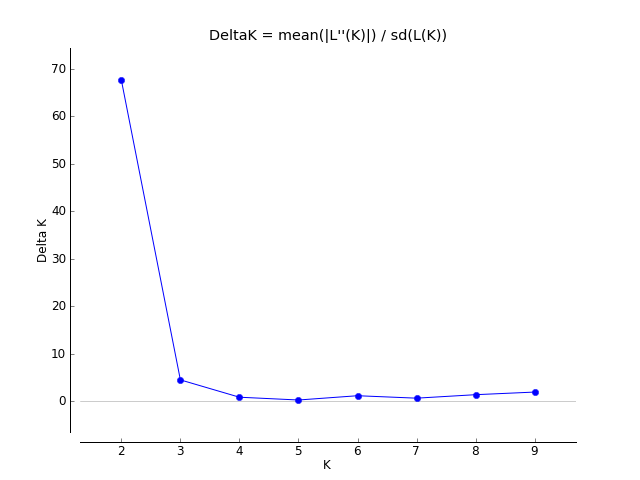

**Supplemental Figure 1.** STRUCTURE HARVESTER graph showing high support for two genetic groups (K = 2, ∆K = 67.68) and weak support for three genetic groups (K = 2, ∆K = 4.48)

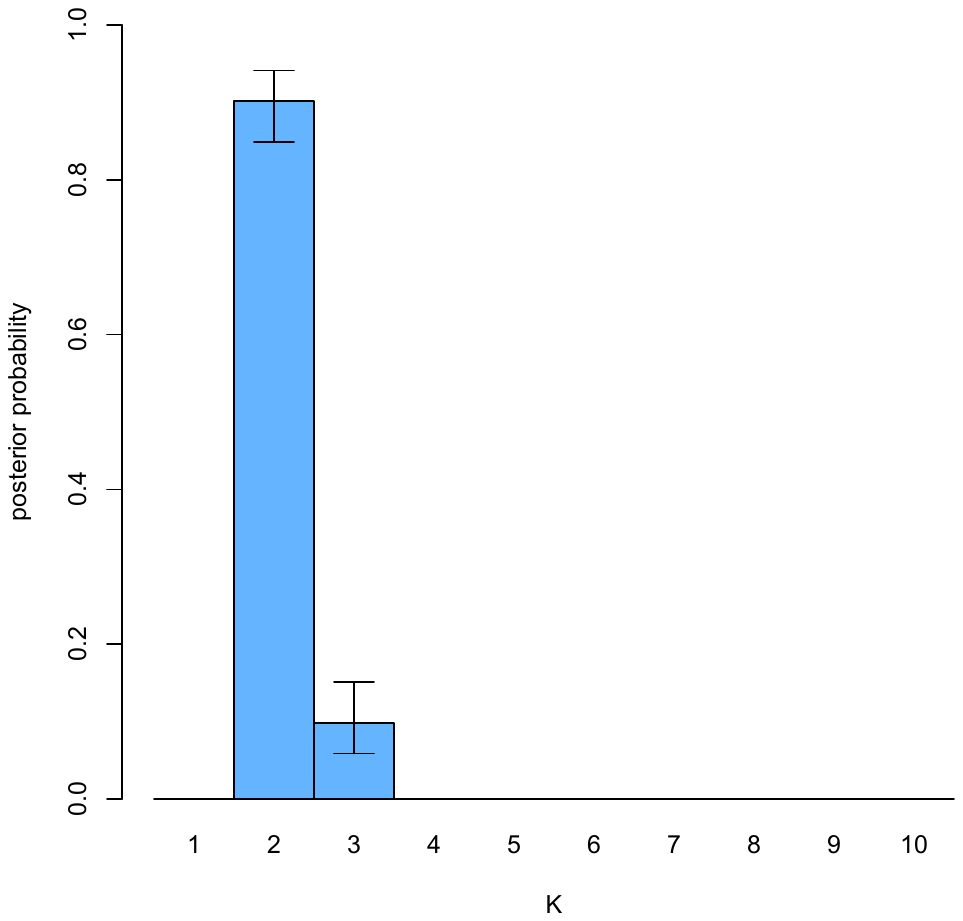

**Supplemental Figure 2.** MavericK 1.0.5 thermodynamic integration evidence estimates normalized to a sum of 1.0.

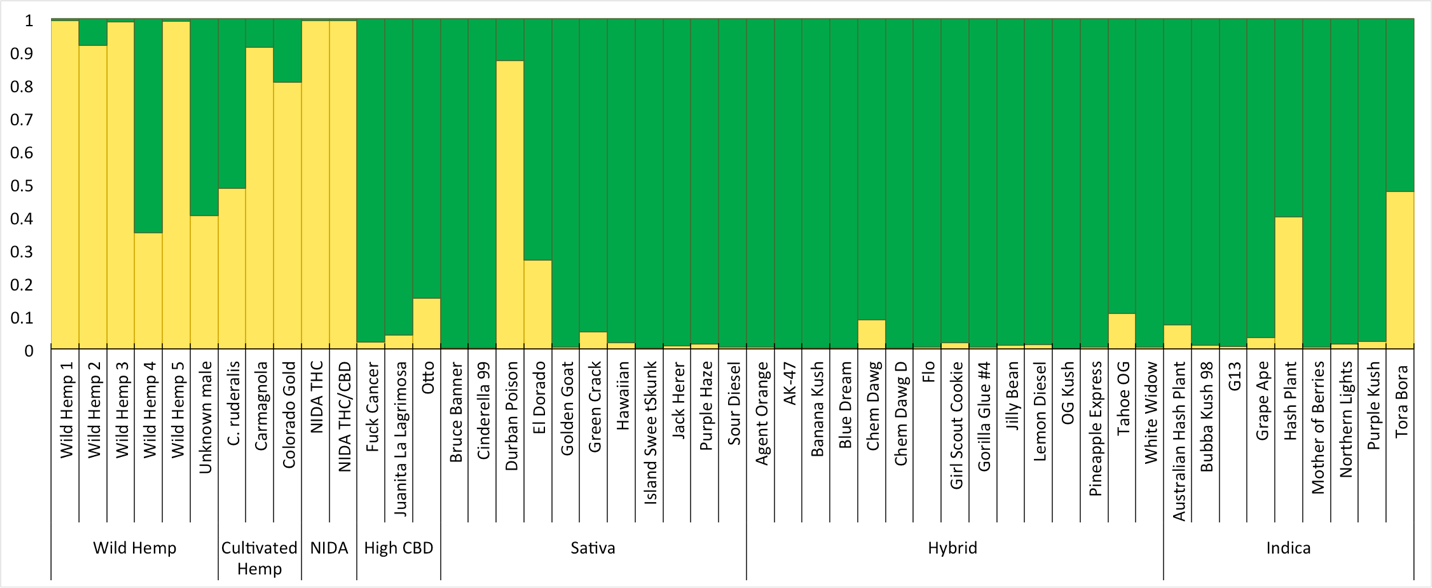

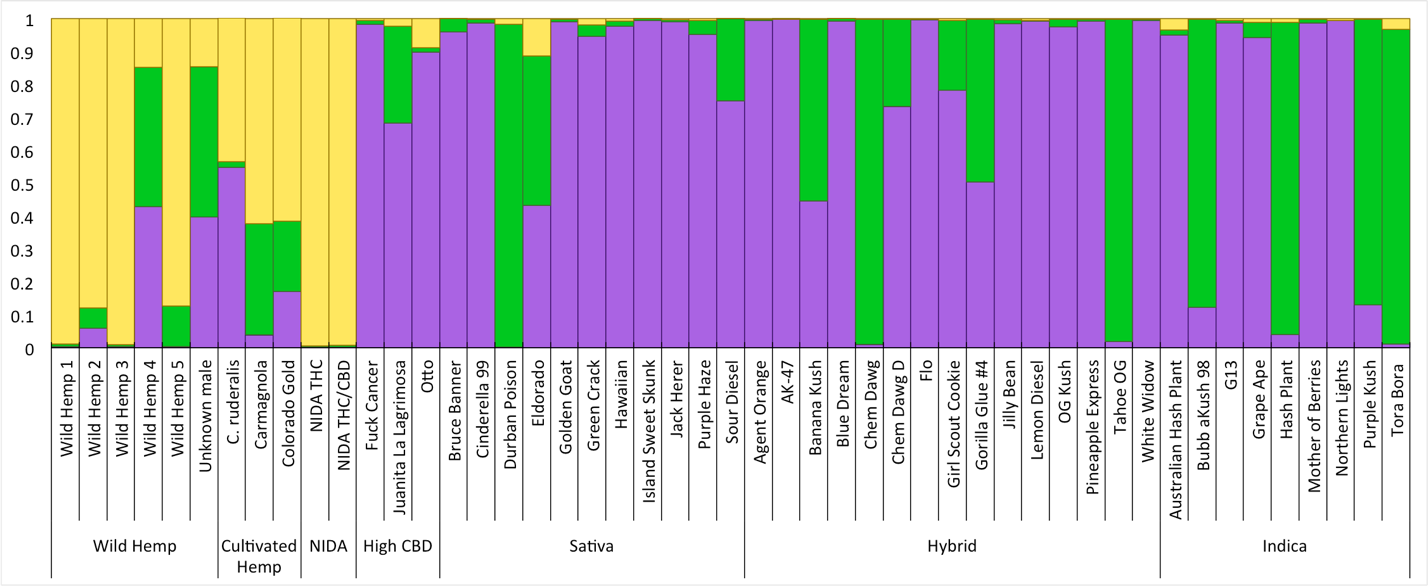

**Supplemental Figure 3.** MavericK 1.0.5 graphs with the proportion of ancestry is represented by two (yellow and green) or three (yellow, green and purple) colors, and each bar represents on individual.

| **Supplemental Table 1**. Sample names, ID code, accession number/ strain name, and the suppliers name and location | | | | | |
| --- | --- | --- | --- | --- | --- |
| **Name** | **ID Code** | **Accession/ Strain Name** | **Supplier Origin** | **City** | **State** |
| Wild Hemp 1 | 1019 | Hemp 1019 | DBG Herbarium | Denver | Colorado |
| Wild Hemp 2 | 24845 | Hemp 24845 Male | DBG Herbarium | Denver | Colorado |
| Wild Hemp 3 | 25572 | Hemp 25572 Male | DBG Herbarium | Denver | Colorado |
| Wild Hemp 4 | 22831M | Hemp 22831 Male | UNC Herbarium | Greeley | Colorado |
| Wild Hemp 5 | 28381M | Hemp 28381 Male | DBG Herbarium | Denver | Colorado |
| Wild Hemp 6 | UnkM | Hemp Unknown Male | Cannabis Genomic Research Initiative | Boulder | Colorado |
| Wild Hemp 7 | Cara#2_4 | Hemp Cara#2 | Cannabis Genomic Research Initiative | Boulder | Colorado |
| Wild Hemp 8 | Carm | Hemp Carmagnola | Colorado Seed, Karen Kershner | Colorado Springs | Colorado |
| Wild Hemp 9 | CoGo | Hemp Colorado Gold | Colorado Seed, Karen Kershner | Colorado Springs | Colorado |
| NIDA THC | NIDA_THC | NIDA High THC | Univ. of Misssissippi | Mississippi | Colorado |
| NIDA THC/CBD | NIDA_THC-CBD | NIDA THC/CBD | Univ. of Mississippi | Mississippi | Colorado |
| Otto (High CBD) | Otto1 | Otto (High CBD) | Centennial Seeds | Colorado Springs | Colorado |
| Juanita La Lagrimosa | JLL_2 | Juanita La Lagrimosa (High CBD) | Nature's Herbs and Wellness | Garden City | Colorado |
| Fuck Cancer | FuCa | Fuck Cancer | Matt Kahl | Colorado Springs | Colorado |
| Durban Poison | DuPo_19 | Durban Poison | The Kind Room | Denver | Colorado |
| El Dorado | ElDo_1 | El Dorado | Smokey's 420 | Garden City | Colorado |
| Hawaiian | Hawa_9 | Hawaiian | Best Colorado Meds | Fort Collins | Colorado |
| Sour Diesel | SoDi_2a | Sour Diesel | Nature's Herbs and Wellness | Garden City | Colorado |
| Island Sweet Skunk | ISS_1 | Island Sweet Skunk | Smokey's 420 | Garden City | Colorado |
| Agent Orange | AgOr_1 | Agent Orange | Nature's Herbs and Wellness | Garden City | Colorado |
| Cinderella 99 | Cin99_1 | Cinderella 99 | Smokey's 420 | Garden City | Colorado |
| AK-47 | AK47_21 | AK-47 | Herbal Alternative | Denver | Colorado |
| Gorilla Glue #4 | GoGl#4_20 | Gorilla Glue #4 | Colorado Wellness | Denver | Colorado |
| Golden Goat | GoGo_19 | Golden Goat | The Kind Room | Denver | Colorado |
| Green Crack | GrCr_2b | Green Crack | Nature's Herbs and Wellness | Garden City | Colorado |
| Bruce Banner | BrBa_19 | Bruce Banner | The Kind Room | Denver | Colorado |
| Flo | Flo_9 | Flo | Best Colorado Meds | Fort Collins | Colorado |
| Jilly Bean | JiBe_1 | Jilly Bean | Smokey's 420 | Garden City | Colorado |
| Pineapple Express | PiEx_2 | Pineapple Express | Nature's Herbs and Wellness | Garden City | Colorado |
| Purple Haze | PuHa_22 | Purple Haze | Lucy Sky | Denver | Colorado |
| White Widow | WhWi_1 | White Widow | Smokey's 420 | Garden City | Colorado |
| Jack Herer | JaHe_12 | Jack Herer | The Milkman | San Luis Obispo | California |
| OG Kush | OGKu_21 | OG Kush | Herbal Alternative | Denver | Colorado |
| Blue Dream | BlDr_19 | Blue Dream | The Kind Room | Denver | Colorado |
| Tahoe OG | TaOG_11 | Tahoe OG | KindCare | Fort Collins | Colorado |
| Chem Dawg | ChDa_8 | Chem Dawg | The Station | Boulder | Colorado |
| Banana Kush | BaKu_2 | Banana Kush | Nature's Herbs and Wellness | Garden City | Colorado |
| Chem Dawg D | ChDaD_19 | Chem Dawg D | The Kind Room | Denver | Colorado |
| Girl Scout Cookie | GSC_14 | Girl Scout Cookie | Day & Night | San Luis Obispo | California |
| G13 | G13_10 | G13 | Infinite Wellness | Fort Collins | Colorado |
| Lemon Diesel | LeDi_2 | Lemon Diesel | Nature's Herbs and Wellness | Garden City | Colorado |
| Hash Plant | HaPl_1 | Hash Plant | Smokey's 420 | Garden City | Colorado |
| Australian Hash Plant | HaPlAu_1 | Australlian Hash Plant | Smokey's 420 | Garden City | Colorado |
| Bubba Kush 98 | Bub98_19 | Bubba Kush | The Kind Room | Denver | Colorado |
| Mother of Berries | MoBe_2 | Mother of Berries | Nature's Herbs and Wellness | Garden City | Colorado |
| Northern Lights | NoLi_15 | Northern Lights | CannaExpress | San Luis Obispo | California |
| Grape Ape | GrAp_16 | Grape Ape | Slow Burn | Union Gap | Washington |
| Purple Kush | PuKu_19 | Purple Kush | The Kind Room | Denver | Colorado |
| Torro Bora | ToBo_4 | Torro Bora | Cannabis Genomic Research Initiative | Boulder | Colorado |

**Supplemental Table 2**. Scored microsatellite data for 49 samples for 10 loci. Missing data represented by zeros.

|  | Casa_002 - 1 | Casa_002 - 2 | Casa_6 - 1 | Casa_6 - 2 | Casa_14 - 1 | Casa_14 - 2 | Casa_18 - 1 | Casa_18 - 2 | Casa_22 - 1 | Casa_22 - 2 | Casa_26 - 1 | Casa_26 - 2 | Casa_27 - 1 | Casa_27 - 2 | Casa_28 - 1 | Casa_28 - 2 | Casa_29 - 1 | Casa_29 - 2 | Casa_30 - 1 | Casa_30 - 2 |
| --- | --- | --- | --- | --- | --- | --- | --- | --- | --- | --- | --- | --- | --- | --- | --- | --- | --- | --- | --- | --- |
| Wild Hemp 1 | 276 | 282 | 416 | 422 | 272 | 272 | 0 | 0 | 192 | 196 | 242 | 242 | 184 | 193 | 175 | 190 | 189 | 189 | 265 | 294 |
| Wild Hemp 2 | 282 | 282 | 416 | 416 | 0 | 0 | 206 | 206 | 184 | 192 | 0 | 0 | 181 | 193 | 175 | 175 | 0 | 0 | 291 | 291 |
| Wild Hemp 3 | 276 | 276 | 410 | 416 | 260 | 260 | 0 | 0 | 192 | 192 | 0 | 0 | 184 | 184 | 181 | 181 | 186 | 195 | 294 | 303 |
| Wild Hemp 4 | 300 | 300 | 410 | 416 | 263 | 263 | 0 | 0 | 192 | 192 | 0 | 0 | 184 | 193 | 175 | 199 | 0 | 0 | 303 | 303 |
| Wild Hemp 5 | 276 | 276 | 416 | 416 | 272 | 272 | 212 | 212 | 184 | 184 | 194 | 194 | 193 | 196 | 190 | 190 | 186 | 186 | 267 | 267 |
| Wild Hemp 6 | 300 | 300 | 416 | 416 | 281 | 281 | 209 | 209 | 184 | 192 | 197 | 239 | 0 | 0 | 175 | 178 | 0 | 0 | 261 | 261 |
| Wild Hemp 7 | 276 | 306 | 416 | 416 | 0 | 0 | 206 | 212 | 184 | 192 | 206 | 206 | 184 | 184 | 178 | 181 | 183 | 186 | 303 | 303 |
| Wild Hemp 8 | 276 | 300 | 416 | 416 | 281 | 281 | 203 | 212 | 192 | 192 | 197 | 197 | 184 | 199 | 178 | 178 | 186 | 195 | 261 | 261 |
| Wild Hemp 9 | 276 | 276 | 410 | 422 | 0 | 0 | 209 | 218 | 184 | 188 | 254 | 254 | 184 | 190 | 193 | 193 | 192 | 195 | 0 | 0 |
| NIDA THC | 276 | 276 | 416 | 416 | 272 | 272 | 191 | 215 | 192 | 192 | 215 | 215 | 184 | 193 | 169 | 169 | 189 | 192 | 265 | 326 |
| NIDA THC/CBD | 276 | 294 | 416 | 416 | 272 | 272 | 191 | 215 | 192 | 192 | 215 | 215 | 184 | 193 | 169 | 169 | 192 | 192 | 326 | 326 |
| Otto (High CBD) | 300 | 300 | 416 | 416 | 260 | 272 | 212 | 221 | 192 | 192 | 206 | 242 | 184 | 184 | 172 | 175 | 183 | 183 | 309 | 309 |
| Juanita La Lagrimosa | 300 | 312 | 416 | 416 | 296 | 296 | 188 | 188 | 192 | 192 | 206 | 206 | 184 | 184 | 0 | 0 | 192 | 192 | 303 | 303 |
| Fuck Cancer | 324 | 324 | 416 | 416 | 275 | 284 | 188 | 221 | 184 | 192 | 206 | 206 | 184 | 184 | 172 | 175 | 195 | 195 | 300 | 300 |
| Durban Poison | 300 | 324 | 416 | 422 | 272 | 275 | 188 | 188 | 188 | 196 | 188 | 188 | 196 | 208 | 190 | 190 | 186 | 195 | 269 | 275 |
| El Dorado | 300 | 300 | 416 | 416 | 272 | 281 | 188 | 188 | 192 | 192 | 206 | 206 | 184 | 193 | 175 | 190 | 183 | 192 | 0 | 0 |
| Hawaiian | 324 | 324 | 416 | 416 | 284 | 287 | 206 | 206 | 192 | 192 | 206 | 206 | 184 | 184 | 178 | 187 | 183 | 192 | 273 | 300 |
| Sour Diesel | 300 | 324 | 422 | 422 | 281 | 281 | 188 | 188 | 192 | 192 | 206 | 206 | 184 | 193 | 190 | 190 | 183 | 192 | 300 | 300 |
| Island Sweet Skunk | 270 | 270 | 410 | 416 | 284 | 297 | 188 | 188 | 184 | 192 | 206 | 206 | 184 | 184 | 175 | 190 | 183 | 192 | 273 | 300 |
| Agent Orange | 276 | 324 | 416 | 422 | 284 | 284 | 212 | 221 | 192 | 192 | 206 | 206 | 184 | 205 | 172 | 190 | 183 | 183 | 300 | 300 |
| Cinderella 99 | 300 | 300 | 410 | 410 | 281 | 284 | 188 | 188 | 192 | 208 | 206 | 206 | 184 | 184 | 175 | 175 | 183 | 183 | 300 | 300 |
| AK-47 | 270 | 324 | 416 | 416 | 284 | 284 | 212 | 221 | 192 | 192 | 206 | 206 | 184 | 193 | 175 | 175 | 183 | 183 | 300 | 300 |
| Gorilla Glue #4 | 300 | 324 | 416 | 416 | 281 | 287 | 188 | 188 | 192 | 192 | 206 | 206 | 184 | 199 | 190 | 190 | 192 | 192 | 273 | 300 |
| Golden Goat | 270 | 270 | 410 | 410 | 284 | 297 | 188 | 212 | 184 | 192 | 206 | 206 | 339 | 339 | 190 | 190 | 183 | 192 | 273 | 300 |
| Green Crack | 270 | 300 | 410 | 416 | 287 | 287 | 182 | 188 | 192 | 192 | 254 | 254 | 184 | 190 | 175 | 202 | 183 | 192 | 300 | 300 |
| Bruce Banner | 300 | 324 | 416 | 416 | 281 | 287 | 188 | 221 | 192 | 192 | 206 | 206 | 184 | 193 | 172 | 190 | 183 | 192 | 300 | 300 |
| Flo | 270 | 270 | 416 | 416 | 279 | 284 | 188 | 188 | 184 | 184 | 206 | 206 | 184 | 205 | 175 | 190 | 183 | 189 | 300 | 300 |
| Jilly Bean | 276 | 276 | 416 | 416 | 284 | 284 | 188 | 188 | 192 | 192 | 206 | 206 | 184 | 184 | 190 | 190 | 183 | 192 | 300 | 300 |
| Pineapple Express | 300 | 300 | 416 | 416 | 287 | 287 | 206 | 212 | 192 | 192 | 206 | 206 | 184 | 184 | 172 | 175 | 192 | 192 | 273 | 300 |
| Purple Haze | 300 | 300 | 416 | 416 | 284 | 287 | 209 | 209 | 184 | 208 | 203 | 206 | 184 | 205 | 172 | 172 | 189 | 192 | 300 | 300 |
| White Widow | 300 | 324 | 416 | 416 | 284 | 284 | 212 | 212 | 184 | 192 | 206 | 206 | 184 | 193 | 175 | 190 | 183 | 192 | 300 | 300 |
| Jack Herer | 270 | 300 | 416 | 422 | 287 | 291 | 185 | 212 | 192 | 192 | 206 | 206 | 184 | 184 | 172 | 172 | 183 | 192 | 265 | 300 |
| OG Kush | 270 | 324 | 416 | 416 | 281 | 287 | 188 | 221 | 192 | 208 | 206 | 206 | 184 | 193 | 172 | 172 | 183 | 192 | 300 | 300 |
| Blue Dream | 270 | 270 | 416 | 416 | 284 | 297 | 188 | 212 | 192 | 208 | 206 | 206 | 184 | 184 | 190 | 190 | 183 | 192 | 300 | 300 |
| Tahoe OG | 276 | 324 | 416 | 416 | 281 | 287 | 188 | 188 | 208 | 208 | 206 | 206 | 196 | 196 | 190 | 190 | 186 | 192 | 275 | 309 |
| Chem Dawg | 300 | 324 | 416 | 416 | 281 | 281 | 188 | 227 | 208 | 208 | 203 | 203 | 193 | 193 | 190 | 190 | 192 | 192 | 275 | 303 |
| Banana Kush | 324 | 324 | 416 | 416 | 281 | 284 | 188 | 227 | 208 | 208 | 206 | 206 | 184 | 196 | 172 | 172 | 183 | 192 | 281 | 281 |
| Chem Dawg D | 270 | 324 | 416 | 416 | 281 | 281 | 188 | 188 | 192 | 192 | 206 | 206 | 184 | 196 | 190 | 190 | 192 | 192 | 300 | 300 |
| Girl Scout Cookie | 276 | 300 | 422 | 422 | 281 | 281 | 188 | 188 | 192 | 192 | 206 | 206 | 184 | 193 | 172 | 172 | 183 | 192 | 265 | 265 |
| G13 | 270 | 270 | 416 | 416 | 279 | 284 | 188 | 227 | 184 | 184 | 206 | 206 | 184 | 205 | 187 | 190 | 183 | 189 | 281 | 303 |
| Lemon Diesel | 276 | 276 | 422 | 422 | 284 | 284 | 212 | 212 | 192 | 192 | 206 | 206 | 184 | 184 | 172 | 175 | 183 | 183 | 300 | 300 |
| Hash Plant | 300 | 300 | 416 | 416 | 272 | 284 | 188 | 188 | 192 | 208 | 188 | 188 | 193 | 196 | 190 | 202 | 183 | 189 | 252 | 252 |
| Australian Hash Plant | 306 | 306 | 416 | 416 | 287 | 287 | 221 | 221 | 184 | 184 | 206 | 206 | 184 | 193 | 172 | 175 | 183 | 189 | 252 | 261 |
| Bubba Kush 98 | 324 | 324 | 410 | 416 | 281 | 281 | 182 | 188 | 192 | 208 | 206 | 206 | 184 | 196 | 190 | 190 | 192 | 192 | 297 | 297 |
| Mother of Berries | 270 | 300 | 416 | 416 | 284 | 287 | 188 | 212 | 192 | 192 | 203 | 203 | 184 | 184 | 172 | 172 | 189 | 192 | 273 | 300 |
| Northern Lights | 276 | 276 | 416 | 416 | 284 | 284 | 212 | 212 | 192 | 192 | 206 | 206 | 184 | 184 | 175 | 175 | 183 | 183 | 300 | 300 |
| Grape Ape | 300 | 300 | 416 | 416 | 272 | 287 | 188 | 188 | 192 | 192 | 206 | 206 | 184 | 187 | 172 | 196 | 183 | 195 | 300 | 300 |
| Purple Kush | 294 | 300 | 416 | 422 | 281 | 287 | 188 | 188 | 192 | 208 | 206 | 206 | 196 | 199 | 172 | 190 | 192 | 192 | 273 | 273 |
| Torro Bora | 276 | 276 | 416 | 416 | 272 | 287 | 188 | 191 | 208 | 208 | 203 | 206 | 193 | 199 | 190 | 190 | 186 | 192 | 273 | 273 |
